# Distribution-preserved compression of single-cell atlases for privacy-protected data dissemination and novel cell type discovery

**DOI:** 10.1101/2024.11.16.622584

**Authors:** Zhihan Cai, Zhibin Hu, Shuang-Rong Sun, Zexu Wang, Fan Yang, Jiahuai Han, Feng Zeng

## Abstract

We introduce SUREv2, a tool for constructing lightweight, transmittable, and privacy-preserving references from single cell atlases. SUREv2 introduces a compressed data structure that maintain the distribution of cells within these atlases and develops an out-of-reference scoring method for identifying novel cell populations. This user-friendly tool shall enhance the analysis of single cell datasets by providing a consistent, privacy-focused reference framework.

## Background

Single cell omics technologies provide unprecedented opportunities to discover previously undocumented cell types, referred to here as novel cell types. Identifying these is essential for advancing our understanding of the biological processes and diseases they are involved in. Global initiatives, such as the Human Cell Atlas (HCA)[1] and the Human BioMolecular Atlas Program (HuBMAP)[2], are actively compiling comprehensive single-cell atlas datasets to catalog cell types. Leveraging these atlases, novel cell types can be defined as populations that fall outside the established reference framework. However, constructing a universal reference for systematically identifying novel cell types remains a challenge.

Building a comprehensive reference framework for single-cell data involves several hurdles. First, variations in cell population compositions across datasets complicate integration efforts, and few existing methods address this issue effectively. Second, technical variations across datasets can interfere with mapping new datasets onto reference frameworks. Current methods for batch effect correction are often insufficient for handling differences between reference and user-supplied datasets robustly. Thirdly, the sheer scale of cell atlases makes constructing and updating references time-intensive, especially as new datasets emerge. Additionally, while visualization methods like UMAP (Uniform Manifold Approximation and Projection) are popular, they often fail to accurately position unseen cell types, leading to overlaps with known cell types (Figs. S1 and S2). Finally, there is growing evidence that single-cell data can leak personal information[3], raising privacy concerns that hinder data sharing.

## Results and Discussion

To address these challenges, we developed SUREv2, which compresses reference atlases using a mixture of statistical distributions (Fig. 1a). This compression reduces data volume, supporting cost-effective data sharing while preserving privacy. SUREv2 utilizes a codebook of metacells to represent cellular distributions within the atlas, compressing hundreds of thousands of cells into a minimal set of metacells. Each metacell employs a multivariate model to capture local cell distribution patterns. By sharing only the metacell codebook, data can be reconstructed by generating single-cell count datasets that retain the original distribution patterns. SUREv2 uses asymmetric variational inference[4] to learns both mapping and recovery functions, facilitating the projection of new datasets based on biological signals and mitigating batch effects, as demonstrated in our recent study[5].

**Figure 1.**
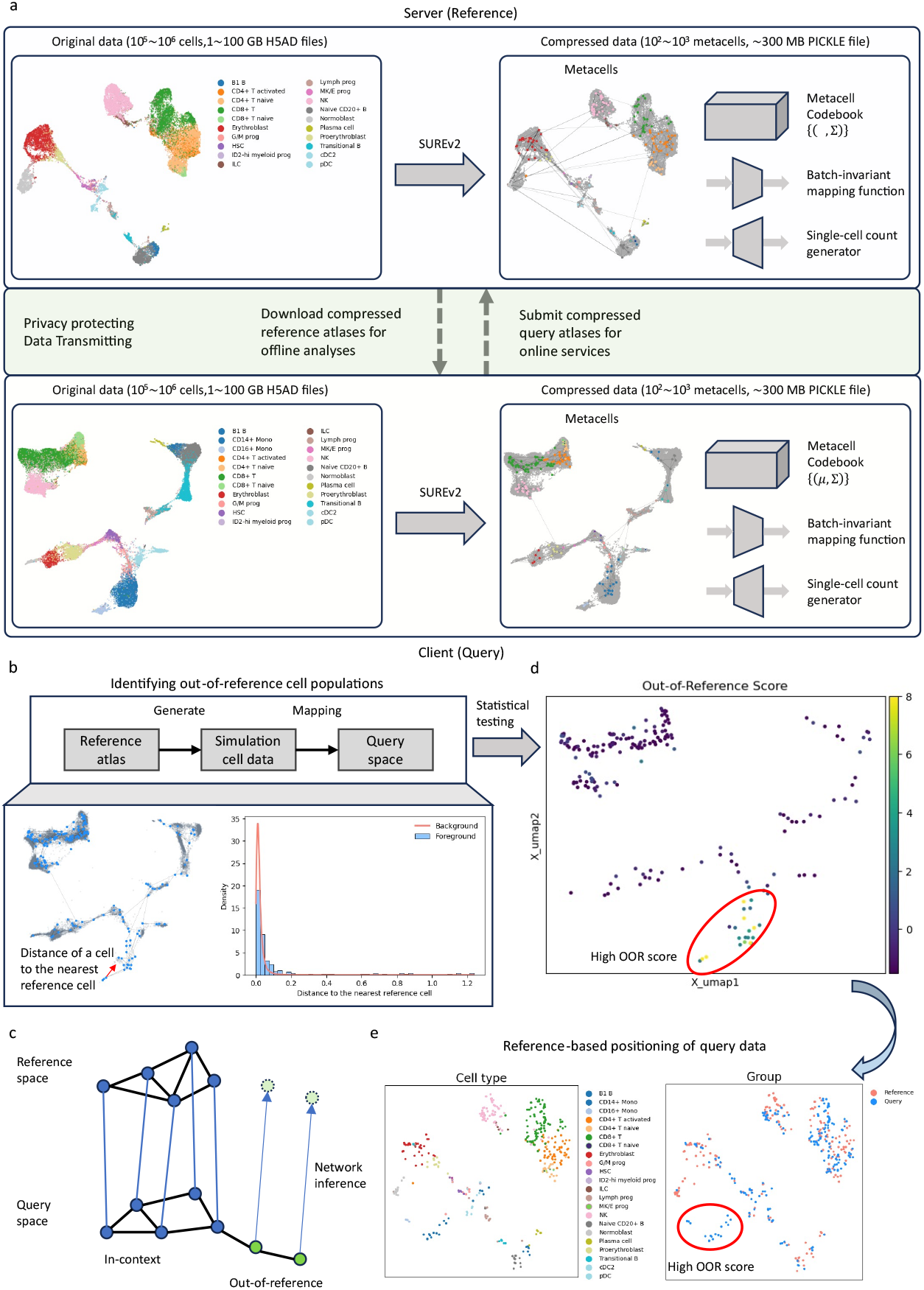
Atlas compression and novel cell type discovery. **a**, Atlases are compressed into a structured form consisting of a metacell codebook, a mapping function, and a reconstruction function. This data structure enables data encoding for shared analysis by both servers and clients. **b**, Diagram of the process for identifying cells that do not align with the reference. **c**, Visualization of query cells mapped within the reference space. **d**, Example of out-of-reference (OOR) scoring for cells within the query dataset. **e**, Visualization of the query dataset using reference-based mapping.

Identifying novel cell types presents additional complexities in reference mapping, as query data may contain cell types absent from the reference framework. While tools like Symphony[6] and scArches[7] have adopted iterative mapping and fine-tuning strategies, they fall short in reliably identifying truly novel cells. SUREv2 introduces a statistical approach to calculate out-of-reference (OOR) scores (Fig. 1b). It first samples cells from the reference and maps them into the query space to establish a background model. This model describes the distance distribution between reference cells and their nearest neighbors. Novel cells are expected to reside farther from the reference cells, while known cells remain proximal. SUREv2 thus scores the novelty of a query cell according to the likelihood of observing its shortest distance to reference cells under the background model. SUREv2 also employs a similarity network to visualize query cells, positioning in-context query cells based on proximity to the nearest reference cells and placing out-of-reference query cells within a network of similar query metacells (Fig. 1c).

SUREv2 enhances the identification of novel cell types within query datasets. To test SUREv2’s capacity for novel cell type identification, we used a dataset from the NeurIPS 2021 Open Problems in Single-cell Analysis[8], splitting it into reference and query sub-datasets. Monocytes (both CD14+ and CD16+ subtypes) were excluded from the reference to simulate novel cell types. SUREv2 successfully identified these monocyte subtypes, assigning accurate novelty scores (Fig. 1d, Fig. S3) and appropriately positioning them in the reference space (Fig. 1e, Fig. S4). By contrast, Symphony failed to distinguish these cells, placing them in overlapping regions with other cell types (Fig. S5), while scArches separated the monocytes but could not differentiate between CD14+ and CD16+ subtypes (Fig. S6).

We assess the limit of atlas compression. Single-cell datasets contain a finite number of distinct cell states, which can be represented in a highly compact form. Using SUREv2, we compressed atlas datasets and evaluated the recovered data using the structural similarity index measure (SSIM) (Fig. 2a, Fig. S7). Our results indicate that: **a)** an atlas can be highly compressed, and **b)** the compression limit is independent of data volume. We tested this with two datasets of varying sizes. When 800 metacells were used for compression, both datasets can be precisely reconstructed, achieving SSIM values greater than 0.85 (Fig. 2b).

**Figure 2.**
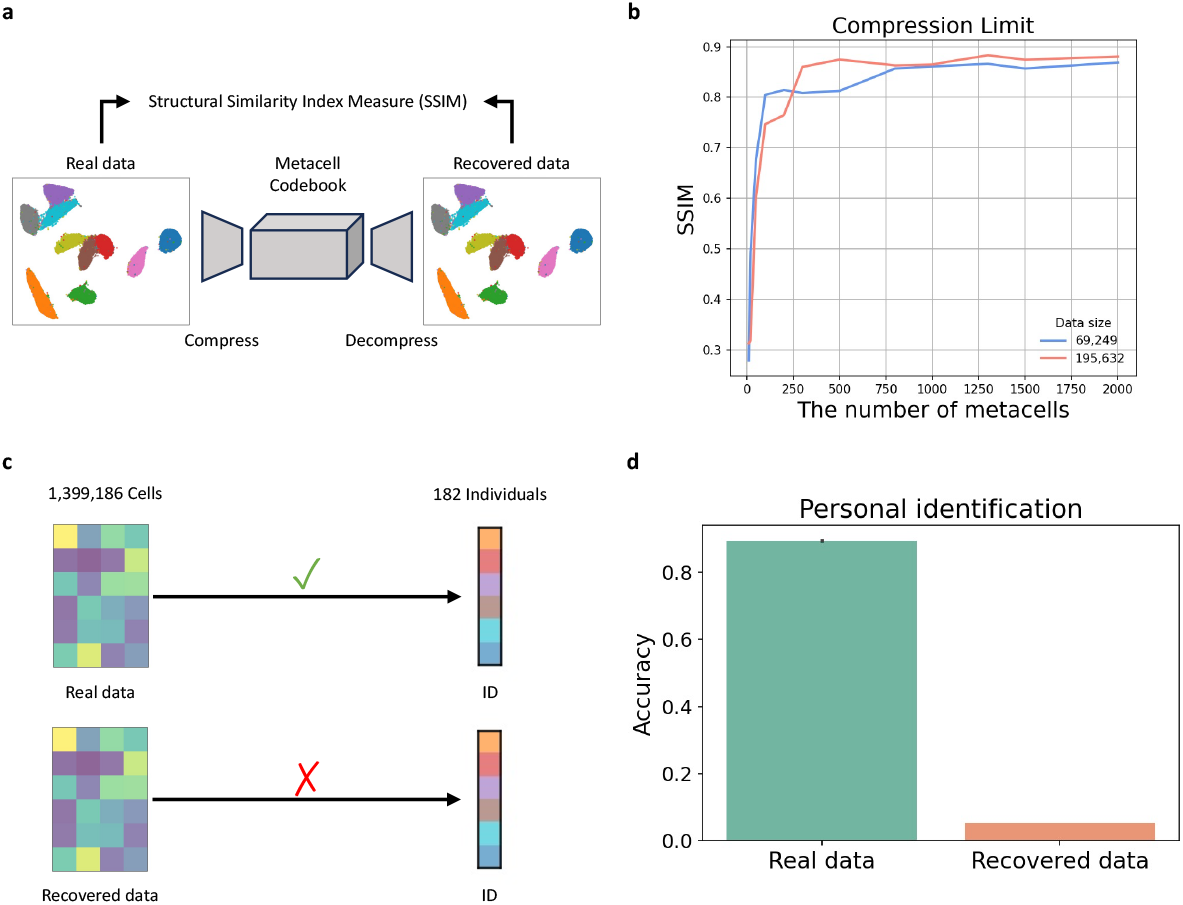
Compression limit and personal identification. **a**, Approach for assessing atlas compression limits. **b**, Compression limit evaluation. **c**, Methodology for evaluating personal identification. **d**, Accuracy results for personal identification.

Compressed atlases can prevent personal identification risks associated with single-cell count data. Recent studies have shown that single-cell data may leak private information, posing risks to public health and impeding data sharing[9,10]. SUREv2’s atlas compression framework provides a potential solution to this issue. To validate this, we applied SUREv2 to a public scRNA-seq dataset containing nearly 1.4 million cells from 182 Asian individuals[11]. Cells from each individual were divided into two halves: one for training a logistic regression classifier and the other for evaluating the accuracy of personal identification (Fig. 2c). The results revealed that the single-cell count data accurately identified individuals, achieving an average accuracy of 0.893 (Fig. 2d). We then conducted the same evaluation on single-cell count data recovered from the SUREv2-compressed atlas. Remarkably, the average accuracy of personal identification significantly decreases to approximately 0.053. These results suggest that SUREv2 can effectively address privacy concerns associated with single-cell atlas sharing.

## Conclusions

In conclusion, SUREv2 provides a metacell-based method for creating compressed cell atlases, which enhance data sharing and enable the discovery of novel cell types. As computational platforms evolve to support single-cell foundation models, compressed atlases like those generated by SUREv2 will be integral to efficient data dissemination and analysis.

## Methods

### SUREv2: Atlas Compression

SUREv2 employs vector quantization (VQ), a classical compression algorithm in signal processing. In VQ, a data point, denoted as *x*, is encoded as a codeword *w*_*i*_ from a codebook *B*. Instead of transmitting the original data, only the codeword index *i* is transmitted. The receiver then recovers the data by retrieving *w*_*i*_ from the codebook:

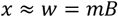

where *m* is a one-hot encoding vector indicating the codeword index.

While traditional VQ is lossy, SUREv2 improves the process by using multivariate distributions rather than discrete codewords. Each multivariate distribution models a distinct cell state or metacell:

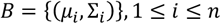

where *μ*_*i*_ and *Σ*_*i*_ are the local and shape parameters of the *i*th metacell.

SUREv2 compresses cell atlases by mapping individual cells into metacells. The dataset is reconstructed through a generation process involving the following steps:

1. **Metacell Selection:** A metacell is selected to generate a cell’s count data;
2. **Latent State Sampling:** The latent state of the cell is sampled from the multivariate distribution of the selected metacell; this latent state is independent of batch effects;
3. **Batch Factor Incorporation:** A batch factor is incorporated to adjust the latent state for batch-specific effects;
4. **Count Data Generation:** The cell’s count data is generated from the negative-binomial model, parametrized by this adjusted latent state.

Let *m, z, b*, and *x* denote the selected metacell, latent state, batch factor, and generated count data, respectively. This process can be summarized as:

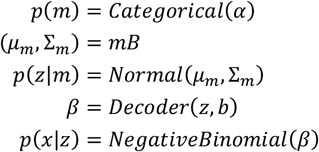

where *α* is the hyperparameter of the distribution over metacells, and *Decoder*(·) is a deep neural network.

SUREv2 learns the metacell codebook and the reconstruction function through an asymmetric variational inference process:

1. **Batch-Free Latent State Inference:** The batch-free latent state of each cell is inferred from its gene expression;
2. **Metacell Assignment:** The cell’s metacell assignment is inferred from this batch-free latent state.

This variational model is represented as:

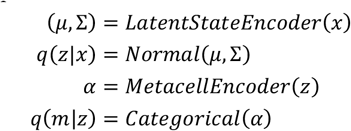

where *LatentStateEncoder*(·) and *MetacellEncoder*(·) are deep neural networks.

The objective for the asymmetric variation inference can be summarized as:

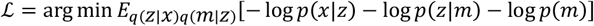

SUREv2 applies the Adam optimization algorithm to the above objective to learn the parameters of both encoders and decoders as well as the metacell codebook. SUREv2 is built using Pytorch and Pyro (a probabilistic programming language).

### Out-of-Reference Scoring

SUREv2 introduces a novel statistical approach to evaluate whether a query atlas contain novel cell types. The method assumes that known cell types will be proximal to cells documented in the reference atlas, while novel cell types are expected to be farther from reference cells.

The scoring process involves the following step:

1. **Simulation and Mapping:** Simulated cells are generated from the reference atlas and mapped to the query atlas. Since low-dimensional representational algorithms like UMAP or autoencoders may not handle unseen data correctly, direct mapping of query data to the reference atlas could lead to overlap between novel and known cell types. To prevent this issue, SUREv2 instead maps the reference data to the query atlas.
2. **Background Distribution Estimation:** Using kernel density estimation, the background distribution *p*_*ref*_ is estimated for the distances between known cell types and their nearest reference neighbors.
3. **Out-of-Reference Scoring:** The background distribution *p*_*ref*_ is then applied to the query data to assess the likelihood of observing the shortest distance *d* between a query cell and the reference atlas. The out-of-reference (OOR) score is defined as

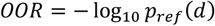

The pseudocode of the above process is given as follows:

| **Out_of_Reference_Score(reference, query, size)** |  |
| --- | --- |
| **IN** | *reference* and *query*: The compressed atlases for the reference and query;<br>*size*: The number of cells will be sampled from the reference atlas. |
| **OUT** | A list of out-of-reference scores for query cells |
| 1. | $x_{ref} \leftarrow \text{codebook\_generate}(\text{reference}, \text{size})$ // Simulate reference cells |
| 2. | $z_{ref} \leftarrow \text{UMAP}_{\text{query}}(x_{ref})$ // Map reference cells to the query atlas |
| 3. | $knn_{ref} \leftarrow \text{NearestNeighbor}(k = 2).fit\_transform(z_{ref})$ // Use KNN to search<br>//for nearest reference<br>// neighbors |
| 4. | $p_{ref} \leftarrow \text{GaussianKDE}(knn_{ref}(z_{ref})[-1])$ // Estimate the background distribution |
| 5. | $OOR_{\text{query}} \leftarrow p_{ref}(knn_{ref}(z_{\text{query}})[1])$ // Compute out-of-reference scores |

### Metacell Similarity Network

A similarity network for metacells yielded from SUREv2 can be constructed based on cell assignments. If two metacells contain a similar set of cells, they are considered similar. In this network, the similarity between metacells *j* and *k* is defined as

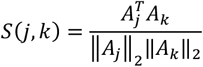

where *A* is the assignment matrix, with rows representing cells and columns representing metacells. SUREv2 uses this similarity matrix S as an adjacency matrix to construct the network of metacells.

### Reference-Based Visualization of Novel Cell Types

SUREv2 supports the visualization of query data within the reference space. The process is as follows:

1. **Positioning In-Context Cells:** Cells with known cell types are positioned by mapping reference cells to the query atlas, building a KNN engine, and finding their neighbors in the reference atlas. The positional shift of the in-context cells from the query space to the reference space is defined as

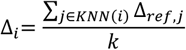

where *Δ*_*ref,j*_ denotes the positional shift of *j*th reference cell from the query space to the reference space.
2. **Positioning Out-of-Reference cells:** After determining positions for in-context cells, out-of-reference cells are placed based on the similarity network. A priority queue maintains unprocessed out-of-reference cells, prioritized by their average distances to in-context cells. The positional shift for an out-of-reference cell is:

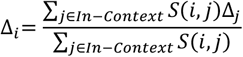

As each out-of-reference cell is positioned, it becomes an in-context cell, and the priority queue is updated.

### Structural Similarity Index Measure

SUREv2 uses the Similarity Index Measure (SSIM) to evaluate the similarity between metacell codebooks and the original cell data. SSIM, commonly applied to assess image similarity, focus luminance, contrast, and structural information, providing a perceptual metric of similarity. A higher SSIM indicates a better preservation of original information after compression.

To adapt SSIM for cell distribution comparison, the dataset is projected into a low-dimensional space (e.g., 2D). This space is divided into grids, and the cell frequency within each grid is estimated to create an image representing the cell distribution. The overall SSIM value is calculated by averaging SSIM values for each cell type.

Let *I*_*k*_ and *J*_*k*_ represent images of the distribution of cell type *k* in original and compressed data, respectively. SSIM is calculated as:

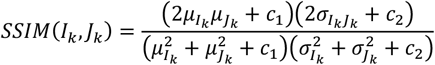

where 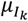 and 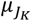 are the means, 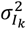 and 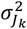 are the variances, and 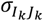 is the covariance. The total SSIM value is defined as:

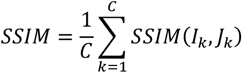

where *C* denotes the number of cell types.

## Acknowledgements

We thank the members of the Han and Zeng group for their comments and help with preparing the manuscript.

## Authors’ contributions

F.Z. and J.H.H conceived the idea and supervised the study. F.Z. and Z.H.C contributed to the code. F.Z., Z.H.C and S.R.S conducted the experiments of novel cell type discovery and personal identification. F.Z., Z.B.H, Z.X.W, and F.Y. conducted the experiment of atlas compression limit. F.Z. wrote the manuscript. F.Z., J.H.H, and F.Y. revised the manuscript.

## Funding

This work is supported in part by the Natural Science Foundation of Xiamen, China (No: 3502Z202373020), National Natural Science Foundation of China (82388201, 61503314), Natural Science Foundation of Fujian Province (2019J01041), the National Key R&D Program of China (2020YFA0803500), the CAMS Innovation Fund for Medical Sciences (2019-I2M-5-062).

### Availability of data and materials

The single-cell atlas datasets used in this study are publicly accessible. The NeurIPS 2021 Open Problem Challenges’ dataset was downloaded from the NCBI GEO depository with the accession number GSE194122. We also involved an additional dataset to evaluate the impact of large-scale data. This dataset, containing 195,632 cells, was download from the CellxGene portal with the link https://cellxgene.cziscience.com/collections/ecb739c5-fe0d-4b48-81c6-217c4d64eec4. For personal identification, the used dataset was also available in the CellxGene portal. Its link is https://cellxgene.cziscience.com/collections/0a839c4b-10d0-4d64-9272-684c49a2c8ba.

SUREv2 are available for academic use on GitHub at https://github.com/ZengFLab/SUREv2.

## Declarations

### Ethics approval and consent to participate

No ethical approval was required for this study

