## Supplemental Figures for "Distribution-preserved compression of single-cell atlases for privacy-protected data dissemination and novel cell type discovery"

### SUPPLEMENTARY FIGURES

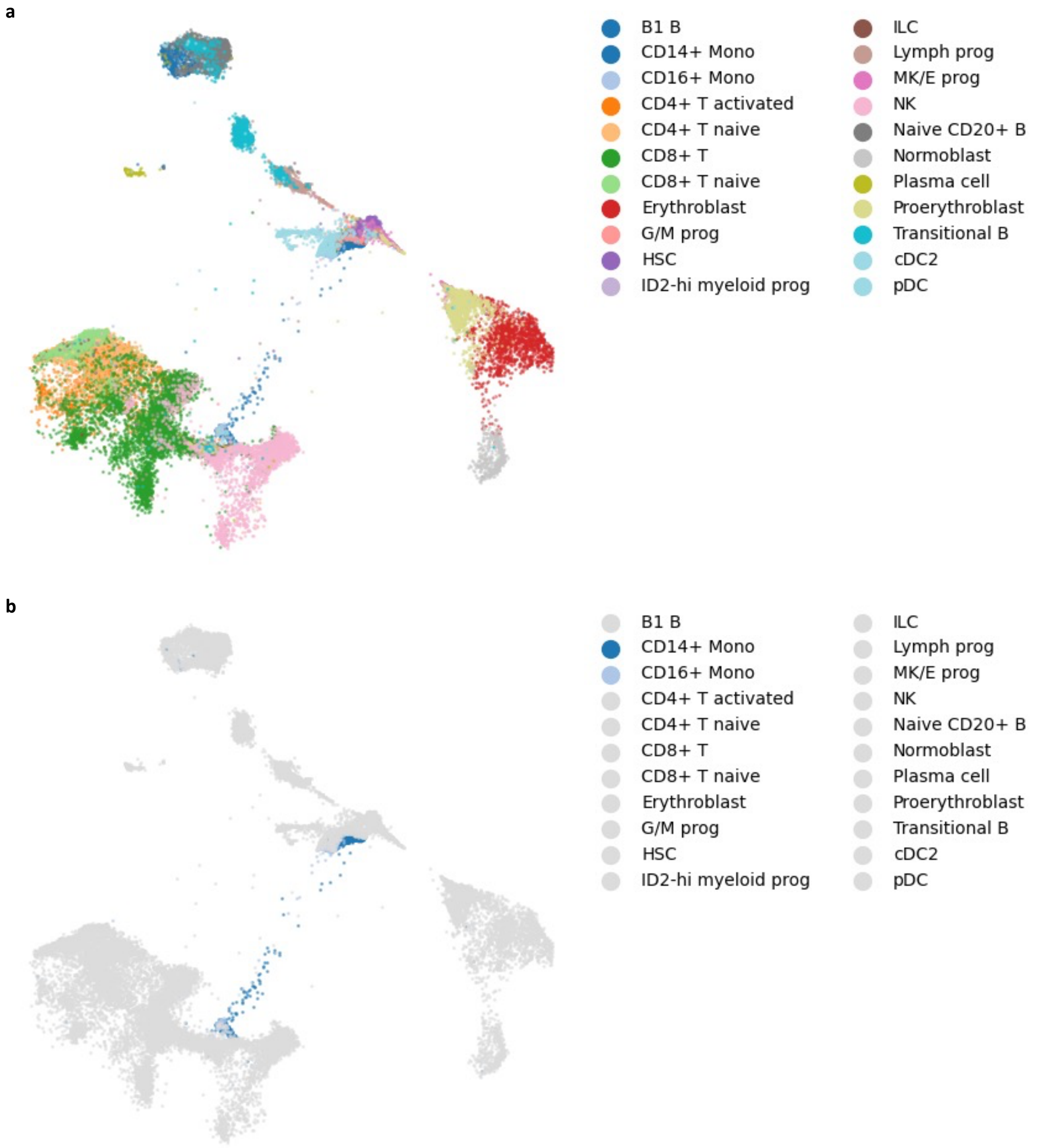


**Supplementary Figure S1. Illustration of UMAP limitations in mapping unseen data.** To highlight UMAP limitations, the NeurIPS 2021 dataset was divided into reference and query subsets, excluding monocytes from the reference subset. After training UMAP on the reference subset, it was applied to the query subset. Monocytes overlapped spatially with existing cell types in the mapping. **a**, Distribution of cell types. **b**, Position of monocytes.

a

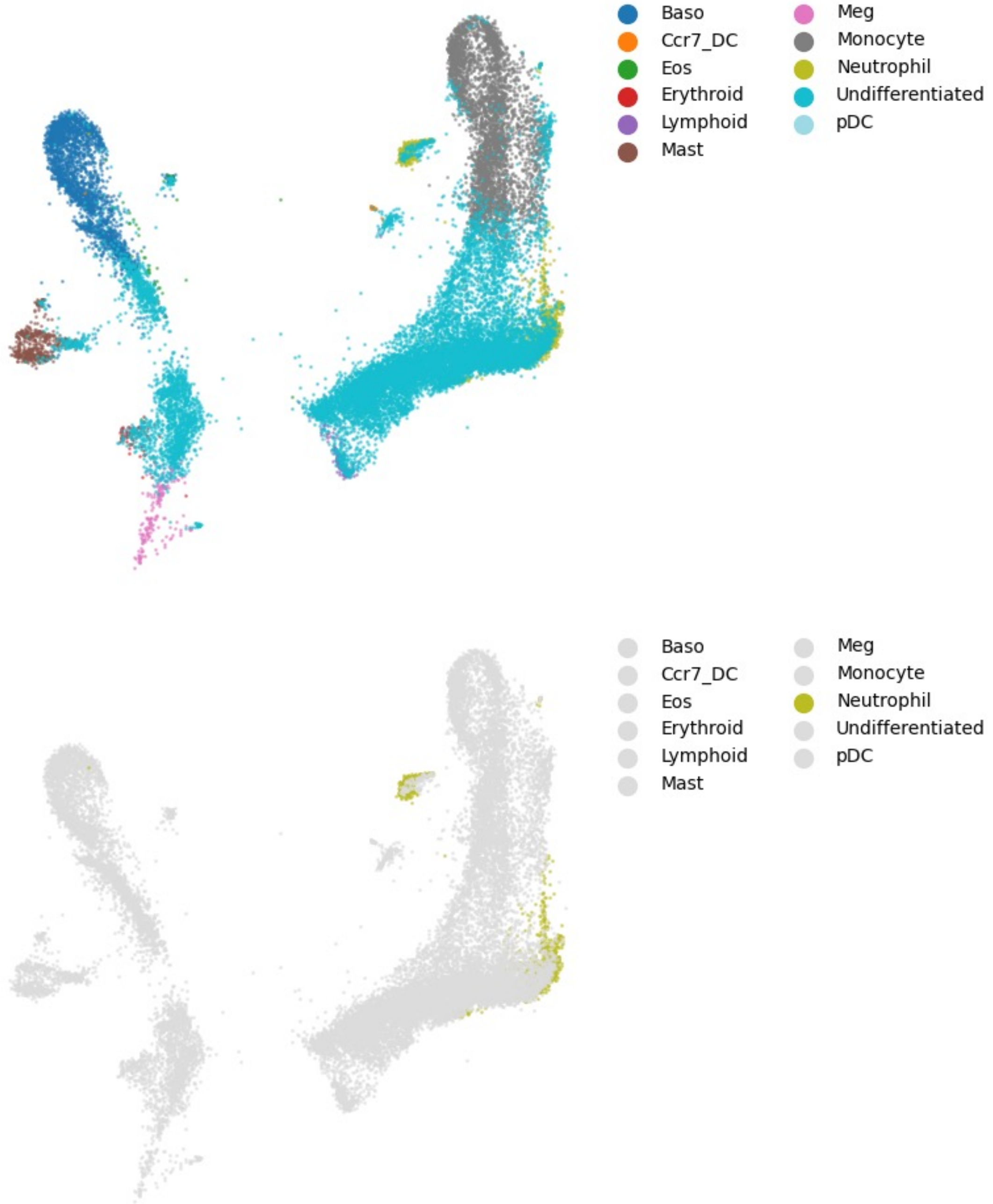


**Supplementary Figure S2. Another example of novel cell type mapping using UMAP.** Using a hematopoiesis dataset, we simulated novel cell type detection by excluding neutrophils from the reference subset. **a**, Distribution of cell types. **b**, Position of neutrophils in the query mapping.

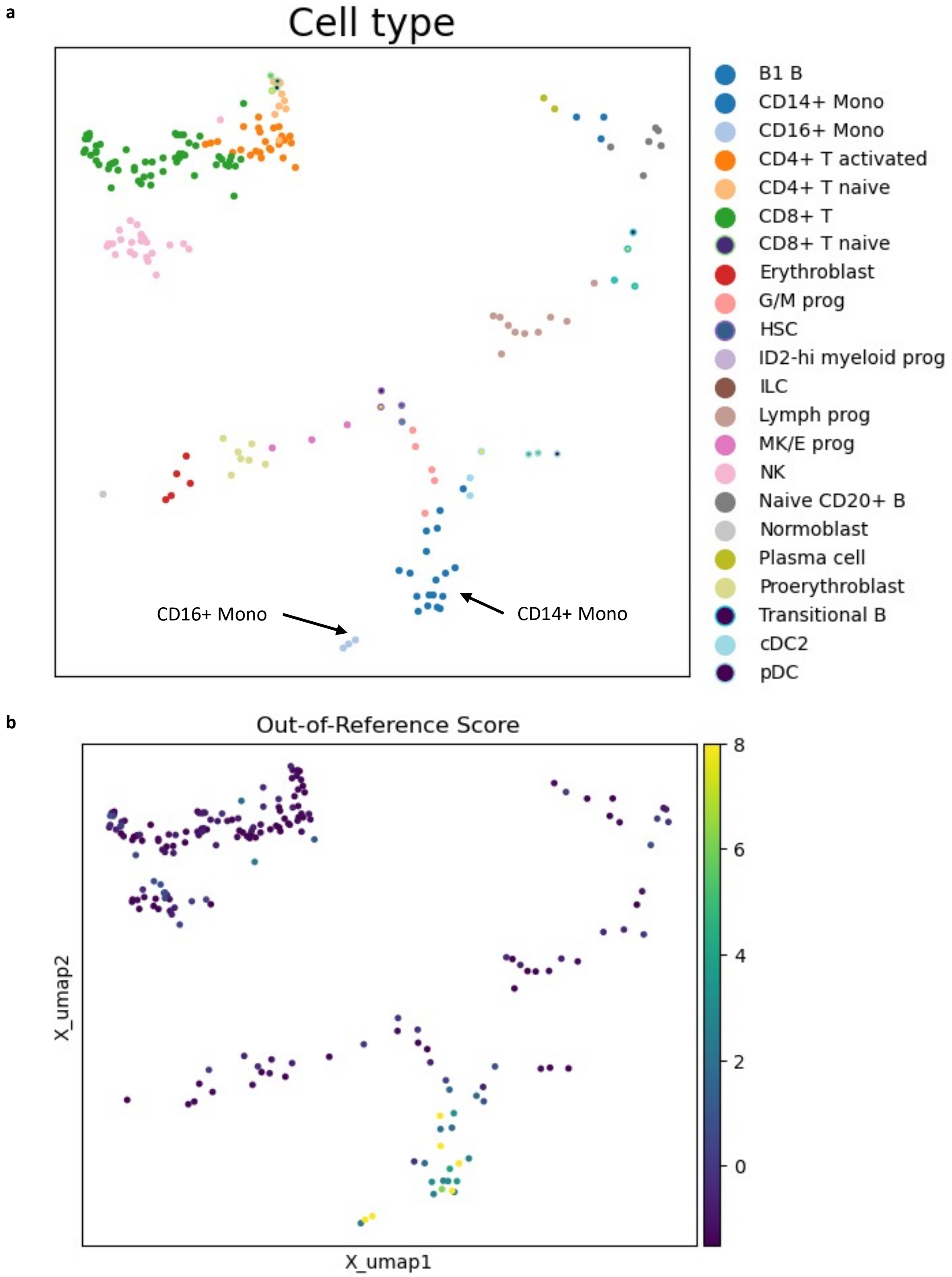


**Supplementary Figure S3. Query atlas and out-of-reference scores from SUREv2. a**, Visualization of query atlas within the query space. **b**, Out-of-reference scores for query metacells as generated by SUREv2.

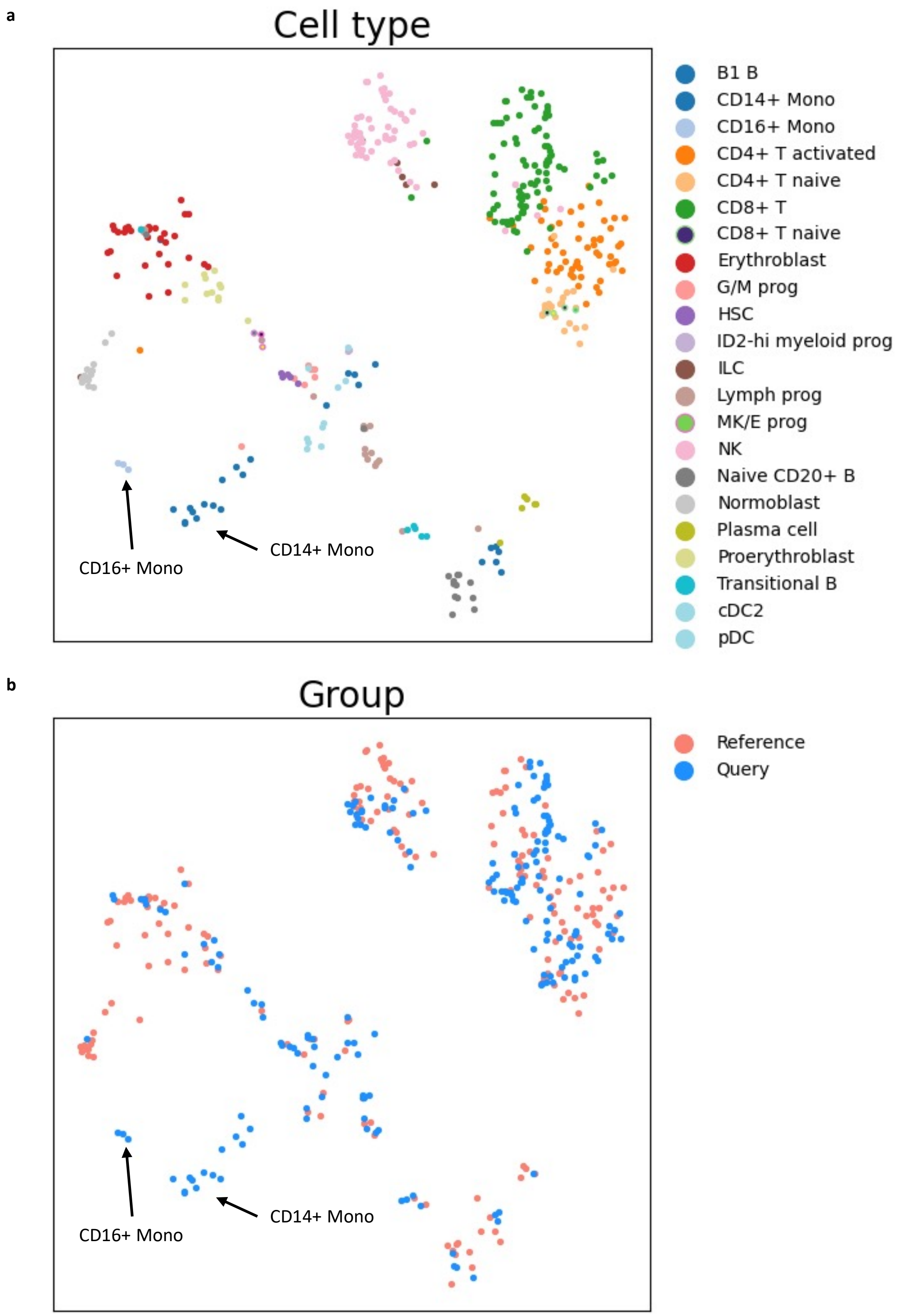


**Supplementary Figure S4. Alignment of query and reference atlases. a**, Distribution of cell types after aligning the query atlas with the reference atlas. **b**, Distribution of distinct groups.

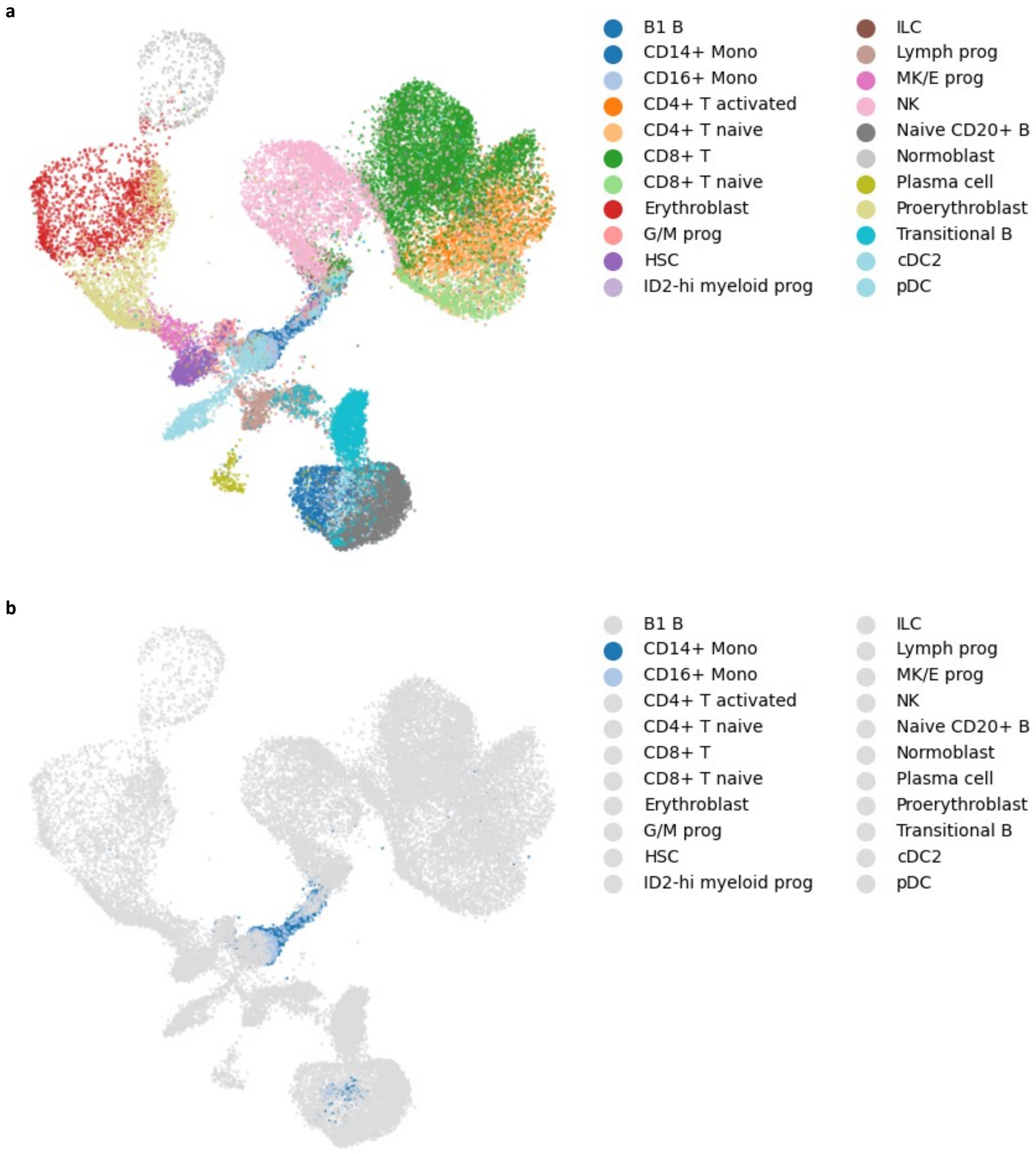


**Supplementary Figure S5. Reference mapping of query data by Symphony. a**, Distribution of cell types following reference mapping. **b**, Distribution of monocytes in the mapped data.

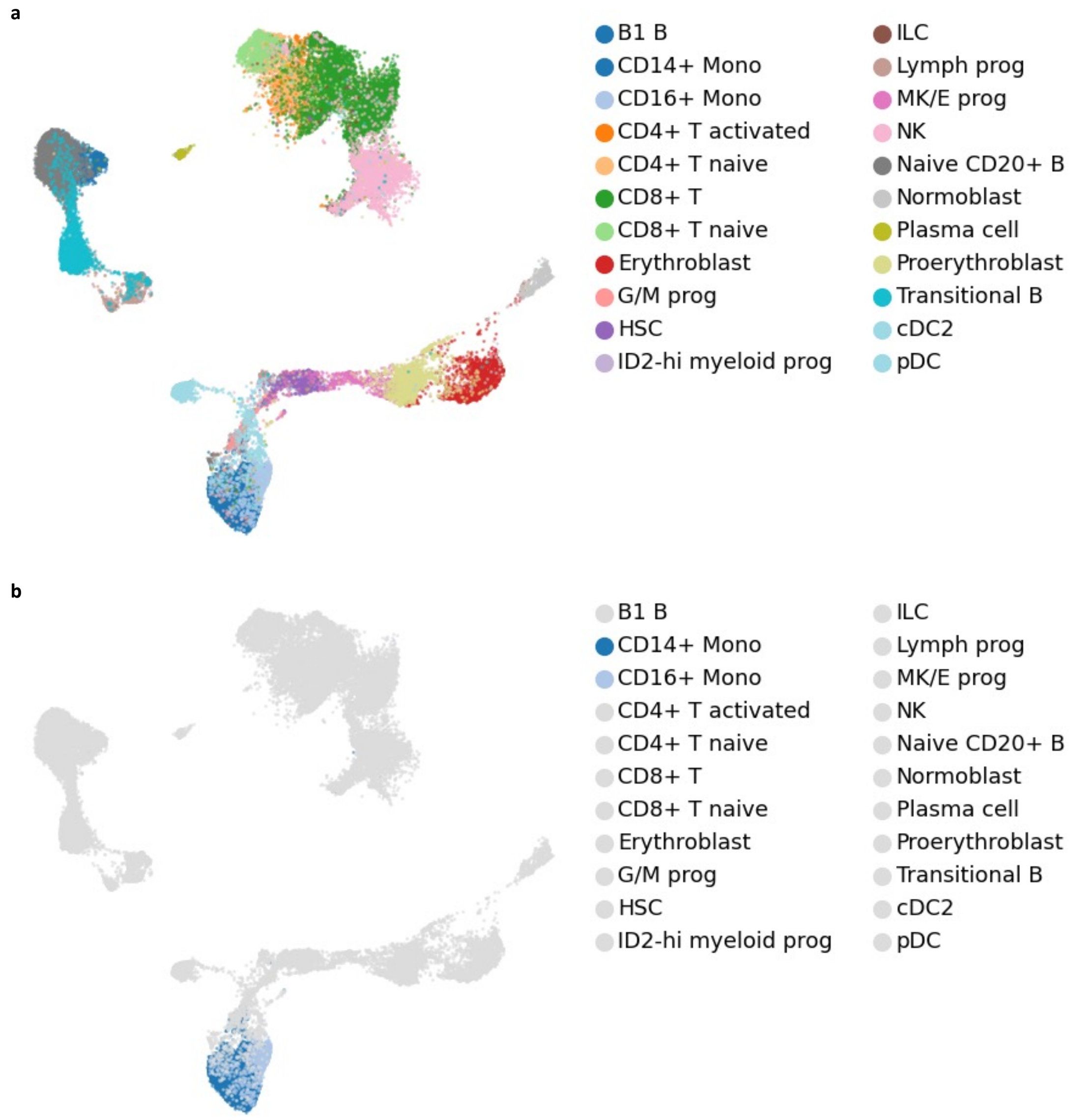


**Supplementary Figure S6. Reference mapping of query data by scArches. a**, Distribution of cell types after applying reference mapping. **b**, Distribution of monocytes in the mapped data.

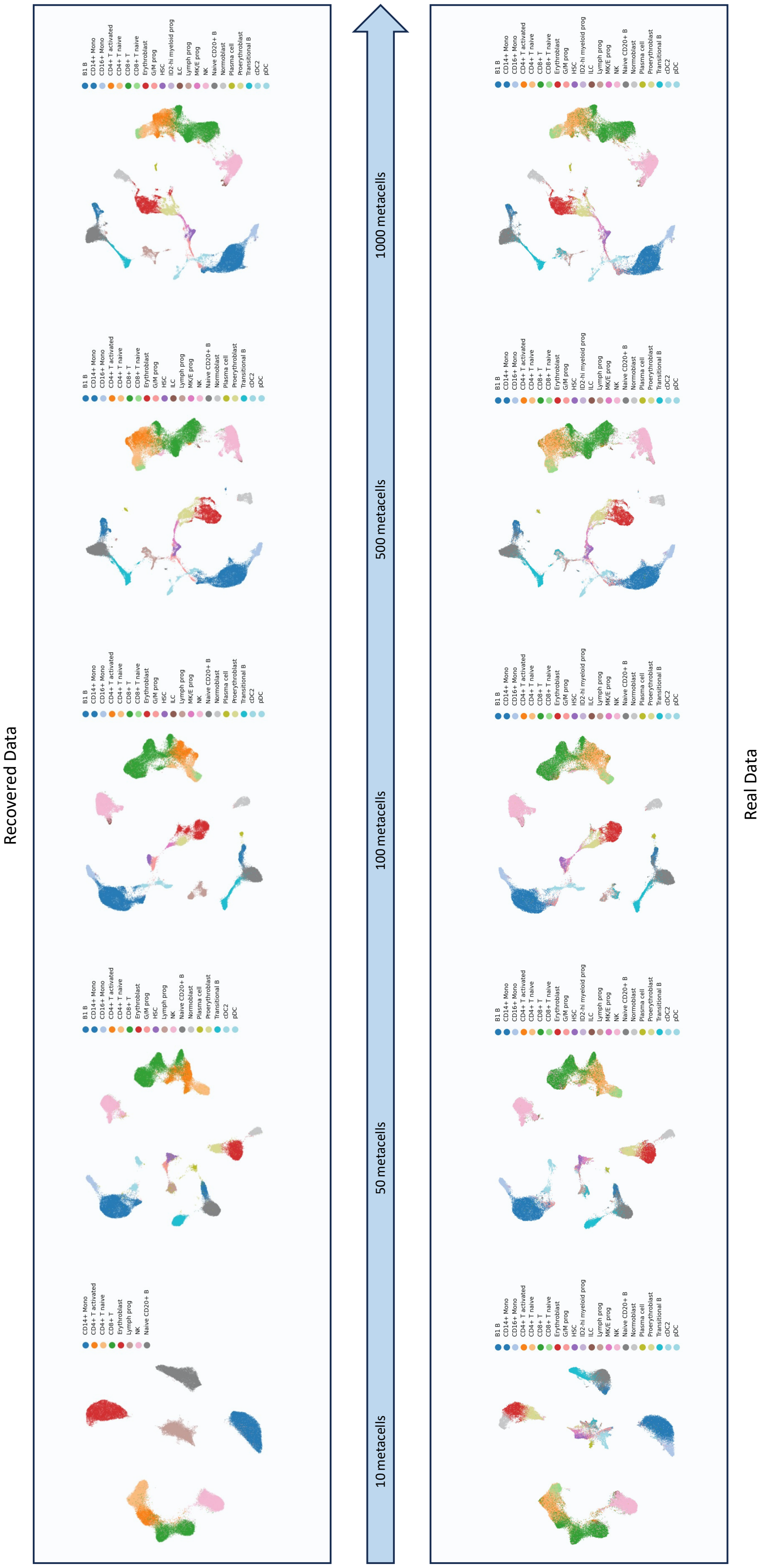


**Supplementary Figure S7. Example of real and recovered data.**
